# T cell co-stimulatory receptor CD28 is a primary target for PD-1–mediated inhibition

**DOI:** 10.1101/086652

**Authors:** Enfu Hui, Jeanne Cheung, Jing Zhu, Xiaolei Su, Marcus J. Taylor, Heidi A. Wallweber, Dibyendu K. Sasmal, Jun Huang, Jeong M. Kim, Ira Mellman, Ronald D. Vale

## Abstract

Programmed death-1 (PD-1) is a co-inhibitory receptor that suppresses T cell activation and is an important cancer immunotherapy target. Upon activation by its ligand PD-L1, PD-1 is thought to suppress signaling through the T cell receptor (TCR). Here, by titrating the strength of PD-1 signaling in both biochemical reconstitution systems and in T cells, we demonstrate that the coreceptor CD28 is strongly preferred over the TCR as a target for dephosphorylation by PD-1- recruited Shp2 phosphatase. We also show that PD-1 colocalizes with the costimulatory receptor CD28 in plasma membrane microclusters but partially segregates from the TCR. These results reveal that PD-1 suppresses T cell function primarily by inactivating CD28 signaling, suggesting that costimulatory pathways may play unexpected roles in regulating effector T cell function and therapeutic responses to anti-PD-L1/PD-1.

## Main Text

T cells become activated through a combination of antigen-specific signals from the T cell receptor (TCR) along with antigen-independent signals from co-signaling receptors. Two sets of co-signaling receptors are expressed on T cell surface: co-stimulatory receptors, which deliver positive signals that are essential for full activation of naïve T cells, and co-inhibitory receptors, which decrease the strength of T cell signaling (*1*). The co-inhibitory receptors serve as “checkpoints” against unrestrained T cell activation and play an important role in maintaining peripheral tolerance as well as immune homeostasis during infection (*2*). One such receptor is PD-1, which binds to two ligands, PD-L1 and PD-L2, which are expressed by a variety of immune and non-immune cells (*3-5*). The expression of PD-L1 is induced by interferon-γ and thus indirectly controlled by T cells which secrete this cytokine upon activation (*4-6*). In addition, T cell activation increases the expression of PD-1 on the T cell itself (*3*). Thus, during chronic viral infection, T cells become progressively “exhausted”, reflecting a homeostatic negative feedback loop due to increased expression of PD-1 and PD-L1(*7-9*). The interaction between PD-1 and its ligands also has been shown to restrain effector T cell activity against human cancers (*10-14*). Antibodies that block the PD-L1/PD-1 axis have exhibited durable clinical benefit in a variety of cancer indications, especially in patients exhibiting evidence of pre-existing anti-cancer immunity by expression of PD-L1 (*15-19*). Interestingly, benefit often correlates with PD-L1 expression by tumor infiltrating immune cells rather than by the tumor cells themselves.

Despite its demonstrated importance in the treatment of human cancer, the mechanism of PD-1-mediated inhibition of T cell function remains poorly understood. Early work demonstrated that binding of PD-1 to PD-L1 causes the phosphorylation of two tyrosines in the PD-1 cytoplasmic domain, presumably via Lck (*20*). Co-immunoprecipitation and co-localization studies in transfected cells suggested that phosphorylated PD-1 then recruits, directly or indirectly, the cytosolic tyrosine phosphatase Shp2, Shp1, and the inhibitory tyrosine kinase Csk (*20, 21*). Defining the direct targets of inhibitory effectors will be critical to understanding the mechanism of anti-PD-L1/PD-1 immunotherapy. However, the downstream targets of PD-1-bound effectors remain poorly understood. Current studies have suggested that PD-1 activation suppresses T cell receptor (TCR) signaling (*21-23*), CD28 costimulatory signaling (*24*), ICOS costimulatory signaling (*25*), or a combination of pathways. Decreased phosphorylation of various signaling molecules, such as Erk, Vav, PLCγ and PI3 kinase (PI3K), has been reported (*21, 24*), but these molecules are common effectors shared by both the TCR and costimulatory pathways and also may not be direct targets of PD-1. Here, we have sought to identify the immediate targets of PD-1 bound phosphatase(s) through a combination of in vitro biochemical reconstitution and cell-based experiments.

We reasoned that the physiological targets of PD-1 should come into close contact with PD-1 sometime during T cell signaling. Therefore, we first sought to map the spatiotemporal relationship between PD-1 and two candidate receptor targets (TCR and CD28) in CD8+ T cells. The TCR and CD28 have been shown to reorganize into submicron-size clusters after binding their ligands (*26*). Using TIRF microscopy and a supported lipid bilayer functionalized with an ovalbumin peptide-MHC class I complex (pMHC; TCR ligand) and B7.1 (CD28 ligand), we found that PD-1 strongly co-localized with the costimulatory receptor CD28 in plasma membrane microclusters (Pearson correlation coefficient (PCC) of 0.89 ± 0.05 (mean ± S.D., n = 17 cells). However, significantly less (*P* < 0.0001) overlap was observed between PD-1 and TCR (PCC of 0.69 ± 0.09) (**Fig. 1, Movie S1**). Strong colocalization of PD-1 and CD28 began from the time of initial cell – bilayer contact (0 sec, **Fig. 1B**) and was sustained until the T cells fully spread (30 sec, **Fig. 1B**). The molecules moved centripetally and eventually became segregated into a canonical bull’s eye pattern with a center TCR island surrounded by CD28 and PD-1, with the latter partially excluded from the TCR rich zone (145 sec, **Fig. 1B**). Some degree of CD28 / PD-1 coclustering was also detected in the absence of pMHC, though the two coreceptors remain largely diffuse without TCR activation (**fig. S1**). As shown previously (*21*), PD-1 clusters also represented sites of Shp2 recruitment to the membrane (**fig. S2**). In the absence of PD-L1 on the bilayer, but with pMHC and B7.1 ligands, PD-1 remained diffusely localized (**fig. S3, Movie S2**), indicating that PD-L1 is required to bring PD-1 and costimulatory receptors into close proximity. Overall, these findings indicate that the costimulatory receptor CD28 strongly co-cluster with PD-1 in the same plasma membrane microdomains in stimulated CD8+ T cells, making CD28 a candidate substrate for dephosphorylation by PD-1– bound phosphatases.

**Figure. 1.**
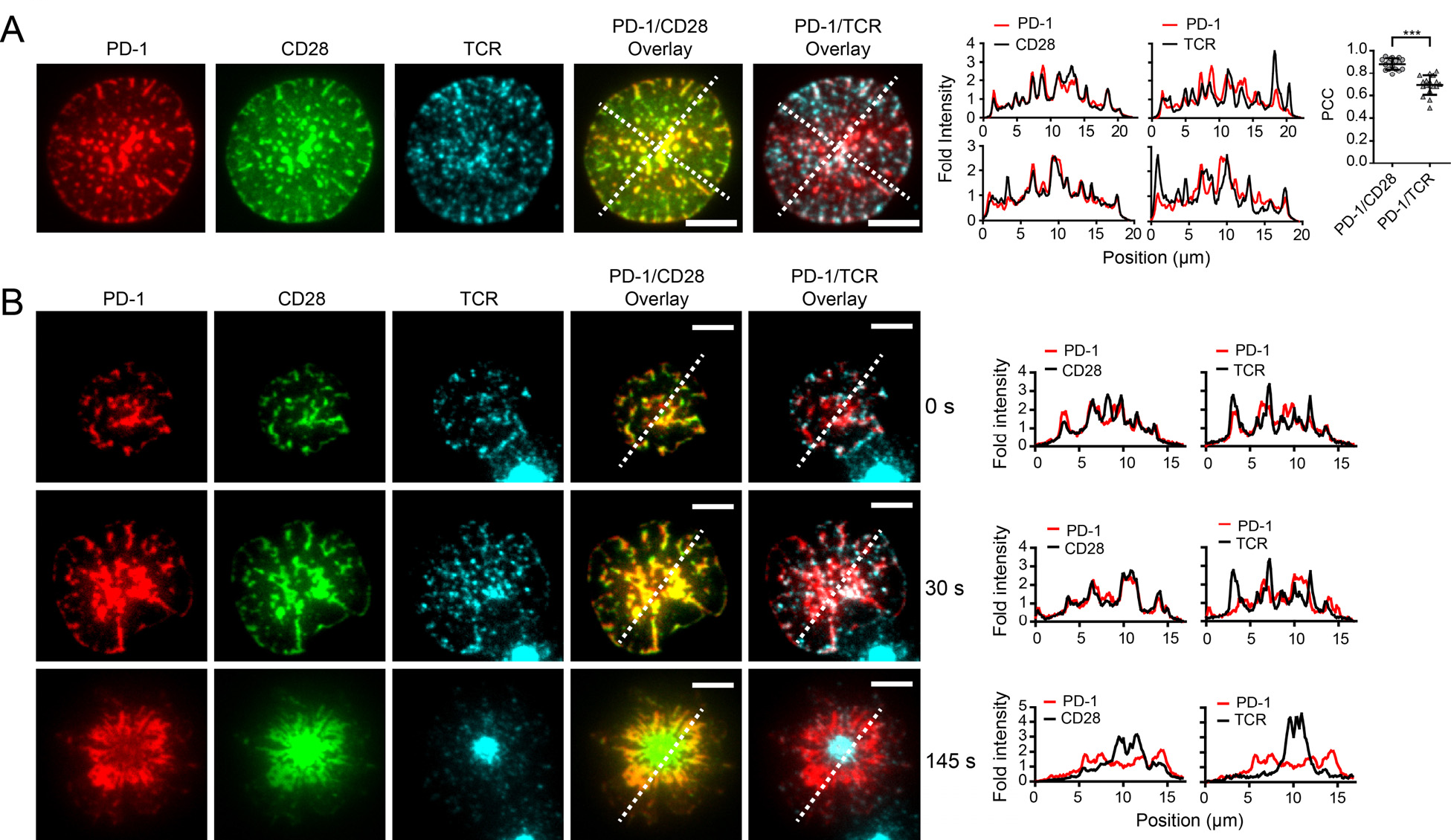
PD-1 coclusters with costimulatory receptor CD28 but partially segregate with TCR. (**A**) (*Left*) Representative TIRF images of PD-1, CD28 and TCR of an OT-I CD8+ T cell 10 sec after landing onto a supported lipid bilayer functionalized with recombinant ligands (100 – 250 molecules per µm^2^), which included peptide-loaded MHC-I (H2Kb), B7.1 (CD28 ligand), and ICAM-1 (integrin LFA1 ligand). Cells were retrovirally transduced with PD-1‒mCherry and CD28‒mGFP and TCR was labeled with an Alexa Fluor647- conjugated anti-TCR antibody (see **Methods)**. (*Middle*) Intensities were calculated from the raw fluorescence intensities along the two diagonal lines in the overlaid images (see **Methods**). PD-1: red; CD28 or TCR: black. (*Right*) Column scattered plot summarizing the Pearson’s correlation coefficient (PCC) values for PD-1/CD28 overlay (0.89 ± 0.05, mean ± S.D) and for PD-1/TCR overlay (0.69 ± 0.09) of 17 fully spread cells, with each dot representing a unique cell. Statistical significance was evaluated by two tailed Student’s t test, *P*<0.0001. (**B**) TIRF images showing the time course of the development of a PD-1/CD28/TCR immunological synapse, starting from initial contact with the supported lipid bilayer (0 sec), to full spreading (30 sec), to a bull eye pattern (145 sec). The histograms from the respective line scan quantifications are shown on the right. Scale bars: 5 μm.

To gain further insight into the substrates affected by activation of PD-1, we turned to a cell-free reconstitution system in which the cytoplasmic domain of the PD-1 was bound to the surface of large unilamellar vesicles (LUVs) that mimic the plasma membrane of T cells (**Fig. 2A**). PD-1 has been shown to co-immunoprecipitate with tyrosine phosphatases Shp2, Shp1, and the Lck-inhibiting kinase Csk from cell lysates (*20*), and contains a structural motif that might recruit the lipid phosphatase SHIP-1 (*27*). Using a FRET-based assay (**Fig. 2A**), we demonstrated that Lck-phosphorylated PD-1 directly binds Shp2 (**Fig. 2B**), but not Shp1, Csk, SHIP-1, or other SH2 proteins tested. Notably, a full titration experiment revealed a 29-fold selectivity of PD-1 towards full length Shp2 over Shp1 (**fig. S4A**) in agreement with qualitative cellular studies (*21*). Surprisingly, the tandem SH2 domains (tSH2) of both Shp1 and Shp2 bind phosphorylated PD-1 with indistinguishable affinities (**fig. S4B**). These data are consistent with a tighter auto-inhibited conformation for Shp1 than Shp2 (*28*), which may decrease Shp1’s affinity for PD-1. Mutation of either tyrosine (Y224 and Y248) in the cytosolic tail of PD-1 led to a partial defect in Shp2 binding (**Fig. 2C** and **fig. S5**). Although Y224 was reported to be dispensable for the ability of PD-1 to co-immunoprecipitate with Shp2 (*29, 30*), our quantitative, direct binding assay shows that both tyrosines in the PD-1 cytosolic domain contribute to Shp2 binding. Collectively, these data suggest that Shp2 is the major effector of PD-1 and that Lck-mediated dual phosphorylation of PD-1 is needed for optimal Shp2 recruitment.

**Figure. 2.**
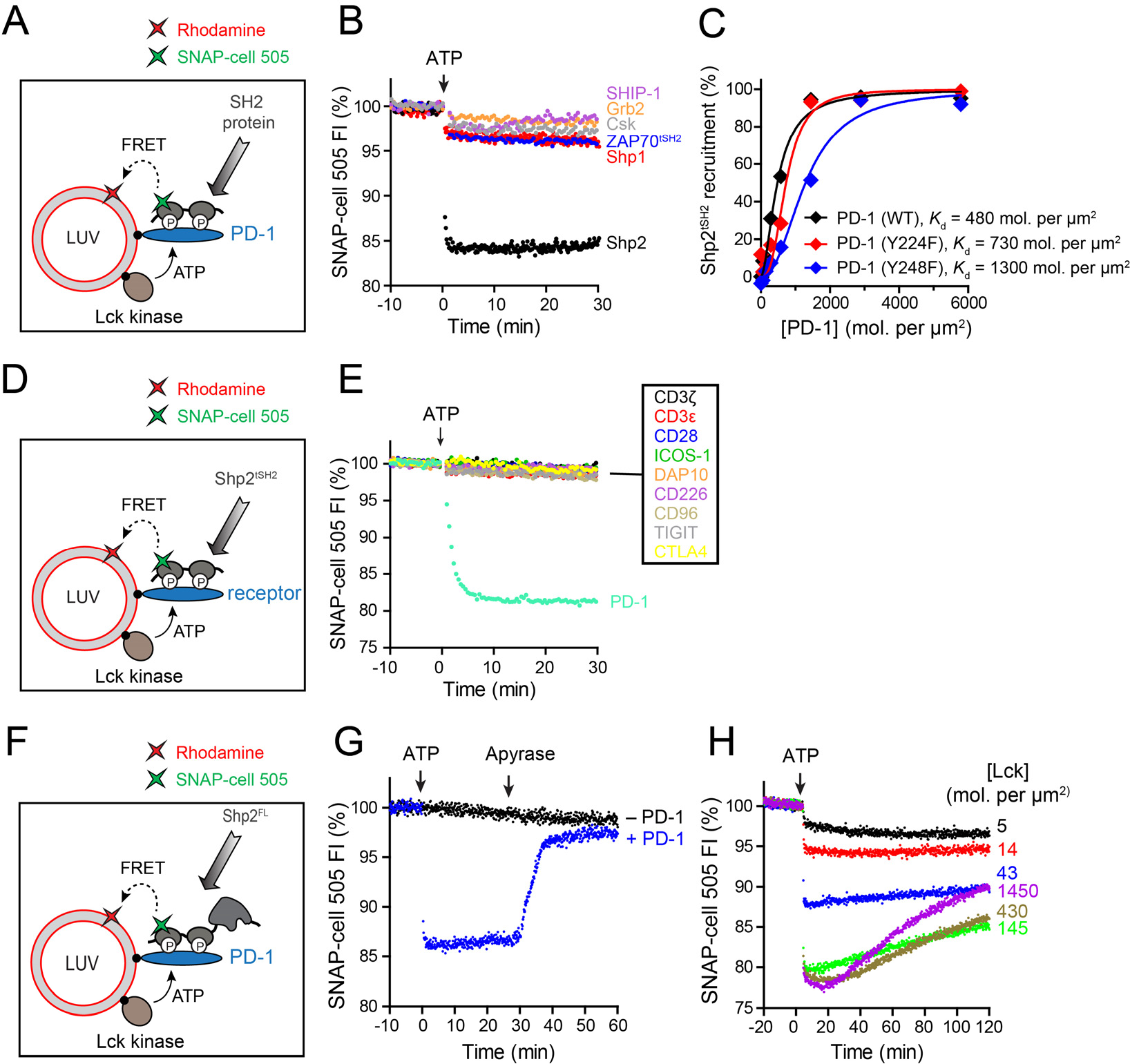
Lck sustains the formation of a highly specific PD-1:Shp2 complex. (**A**) Cartoon depicting a FRET assay for measuring the interaction between a SH2 containing protein and membrane bound PD-1. Rhodamine-PE (energy acceptor) bearing LUVs were reconstituted with purified Lck kinase and the cytosolic domain of PD-1, as described in **Methods**. The SNAP-tag fused SH2 protein of interest was labeled with SNAP-cell 505 (energy acceptor), and presented in the solution. Addition of ATP triggered Lck catalyzed phosphorylation of PD-1 caused the recruitment of certain SH2 proteins to the LUV surface, leading to FRET. (**B**) A comparison of PD-1 binding activities of a panel of SH2 containing proteins, using the FRET assay as described in **A**. Shown are representative time courses of SNAP-505 fluorescence before and after the addition of ATP. Concentrations of components: 870 PD-1 per µm^2^, 21 Lck per µm^2^ and 0.1 μM labeled SH2 protein. (**C**) A comparison of the relative contribution of the two tyrosines of PD-1 in recruiting Shp2. Shown is the degree of Shp2 recruitment against the concentration of LUV-bound PD-1 WT or tyrosine mutant, measured by the FRET assay described in **A**. See **fig. S5** for raw data. (**D**) Cartoon depicting a FRET assay for measuring the ability of a membrane-bound receptor to recruit Shp2. The experiment set up was the same as in **A** except replacing PD-1 with another receptor of interest, and using the tandem SH2 domains of Shp2 (Shp2 ^tSH2^) as a fixed donor bearer. (**E**) A comparison of the Shp2 binding activities of the designated LUV-bound receptors, using the FRET assay shown in **D**. Concentrations: 870 receptor molecules per µm^2^, 21 Lck per µm^2^ and 0.1 μM labeled Shp2^tSH2^. (**F**) Cartoon showing a FRET assay for measuring the localization dynamics of full length Shp2 (Shp2^FL^). Rhodamine-PE (energy acceptor) bearing LUVs were reconstituted with purified Lck kinase and the cytosolic domain of PD-1, as described in **Methods**. SNAP-tag fused Shp2^FL^ was labeled with SNAP-cell 505 (energy acceptor), and presented in the extravesicular solution. **(G)** Time course of the fluorescence of Shp2^FL^*505 in response to the addition of ATP and the ATP scavenger apyrase to the reaction system shown in **F**. A control reaction lacking PD-1 (- PD-1) was run in parallel. **(H)** Time course of the fluorescence of Shp2^FL^*505 showing the dynamics of Shp2 as a function of Lck density. Assay was set up as in **F**.

Using this reconstituted system, we next asked if other signaling receptors besides PD-1 (CD3ζ, CD3ε, CD28, ICOS, DAP10, CD226, CD96, TIGIT, and CTLA4) could recruit Shp2 (**Fig. 2D**). Remarkably, recruitment of Shp2 was not observed for any of these receptors, including two other co-inhibitory molecules TIGIT and CTLA4 (**Fig. 2E**). CTLA4 was reported to co-immunoprecipitate with Shp2 (*31*) and widely believed to suppress T cell signaling at least partly through Shp2 (*32*). Our data suggest that Shp2 does not directly bind CTLA4, and that other proteins are likely required to bridge the two proteins. Overall, our results reveal an unexpected binding specificity of Shp2 for phosphorylated PD-1.

Recruitment of Shp2 to PD-1 suggested that Shp2 might directly dephosphorylate PD-1 to dissemble the PD-1–Shp2 complex, thereby interrupting the negative signaling pathway. To test this idea, we determined the stability of the PD-1–Shp2 complex using a FRET assay (**Fig. 2F**). ATP triggered phosphorylation of PD-1 caused the rapid recruitment of Shp2 (**Fig. 2G**) and activation of its phosphatase activity (**fig. S6**). Termination of the Lck activity by rapid ATP depletion caused a complete dissociation of Shp2 (**Fig. 2G**). This result indicates that the PD-1– Shp2 complex is highly unstable due to the ability of Shp2 to dephosphorylate PD-1 and that continuous Lck kinase activity is required to activate and sustain PD-1–Shp2 mediated inhibitory signaling. Interestingly, a slow spontaneous disassembly of PD-1–Shp2 complex was observed at physiological levels of Lck (**Table S1**), even without the termination of Lck activity (**Fig. 2H**). Collectively, these results demonstrate a positive-negative feedback loop of the Lck–PD-1–Shp2 network in which PD-1–associated Shp2 can override Lck and cause a net kinetic dephosphorylation of PD-1 to dissociate the phosphatase. This feedback regulation would allow the system to quickly reset in the absence of PD-1 ligation or Lck activation.

Having established a highly specific recruitment of Shp2 by PD-1, we turned to identify substrates for dephosphorylation by the PD-1–Shp2 complex. To this end, we reconstituted a diverse set of components involved in the T cell signaling network including: (i) the cytosolic domains of various receptors (PD-1, TCR, CD28, and ICOS (another co-stimulatory receptor (*33*)) on the LUV; (ii) the tyrosine kinases Lck, ZAP70 (a key cytosolic tyrosine kinase which binds to phosphorylated CD3 subunits to propagate the TCR signal (*34*)), and in some experiments the inhibitory kinase Csk (*35*); and (iii) downstream adapter and effector proteins LAT, Gads, SLP76 (*36*) and the regulatory subunit of Type I PI3K (p85α), which is known to be recruited by phosphorylated costimulatory receptors (**fig. S7**) (*37, 38*). All protein components were reconstituted onto LUVs or added in solution at close to their physiological levels (**fig. S8, Table S1**). A reaction cascade consisting of phosphorylation, dephosphorylation, and protein–protein interactions at the membrane surface was triggered by ATP addition. To test the sensitivity of components in this biochemical network to PD-1, we systematically titrated the levels of PD-1 on the LUVs and measured the susceptibility to dephosphorylation of each component in the reactions by phosphotyrosine (pY) western blots **(Fig. 3A)**. This titration also provides insights into how the network responds to the gradual upregulation of PD-1 during T cell development (*39*), activation (*40*), and exhaustion (e.g., in tumors and during chronic virus infections) (*7*).

Strikingly, CD28 but not the TCR or its associated components was found to be the primary target of PD-1–Shp2. As shown in **Fig. 3B,C** (*Left*), CD28 was very efficiently dephosphorylated, with a 50% inhibitory concentration (IC_50_) of ∼96 PD-1 molecules per µm^2^ (**Table S2**). In contrast, PD-1–Shp2 dephosphorylated TCR signaling components to only a minor extent, including the TCR intrinsic signaling subunit CD3ζ, the associated kinase ZAP70, as well as its downstream adaptors LAT and SLP76, whose 50% dephosphorylation occurred at substantially higher PD-1 concentrations (>1000 molecules per µm^2^, **Table S2**). Despite variable degrees of dephosphorylation of the TCR components at high PD-1 levels, a consistently stronger dephosphorylation of CD28 was observed.

Lck, the kinase that phosphorylates TCR, CD28, and PD-1, was the second best target for PD-1–bound Shp2 in the reconstitution system. Both the activating (Y394) and inhibitory (Y505) tyrosines were 50% dephosphorylated at similar levels of PD-1 (400 - 600 molecules per µm^2^). This result, however, suggests a net positive effect of PD-1 on Lck activity, owing to the stronger regulatory effect of the inhibitory tyrosine (*41*). The addition of the Lck inhibiting kinase Csk rendered CD28 as well as TCR signaling components more sensitive to PD-1–Shp2, although CD28 clearly remained the most sensitive PD-1 target (**fig. S9** and **Table S2**). The CD28 preference was robust, as a similar trend was observed at a later time point (**fig. S10**). In contrast to the strong CD28 preference of PD-1/Shp2, the transmembrane phosphatase CD45 efficiently dephosphorylated all of the signaling components tested (**Fig. 3B,C,** *Right*), with only 3-4 fold selectivity on CD28 over CD3ζ and ZAP70 (**Table S2**).

**Figure. 3.**
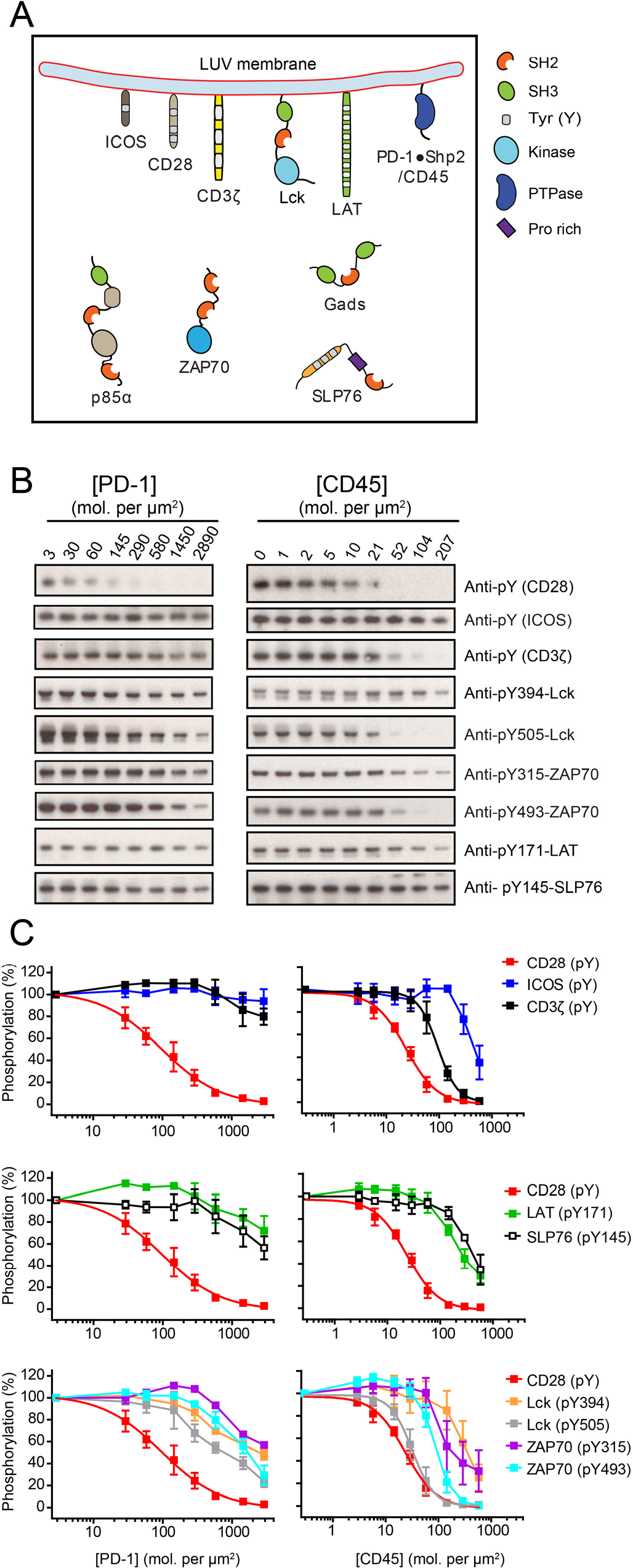
CD28 is uniquely sensitive to PD-1 bound Shp2. (**A**) Cartoon depicting a LUV reconstitution system for assaying the sensitivities of different targets to PD-1:Shp2 or CD45. Purified cytosolic domains of plasma membrane bound receptors (CD3ζ, CD28, PD-1), the adaptor LAT, and the kinase Lck were reconstituted onto LUVs at their physiological molecular densities (**Table S1**). Cytosolic factors (ZAP70, p85α, Gads, SLP76 and Shp2) were presented in the extravesicular solution at their physiological concentrations (**Table S1**). In a parallel experiment, PD-1 and Shp2 were replaced with the cytosolic domain of the tyrosine phosphatase CD45 that was tethered to the LUVs. Addition of ATP triggered a cascade of enzymatic reactions and protein-protein interactions. (**B**) Shp2 containing reactions with increasing [PD-1] or CD45 containing reaction with increasing [CD45] terminated at 30 min, and subjected to SDS-PAGE and phosphotyrosine western blots (WB), as described in **Methods**. (**C**) The optical density of each band in **B** was quantified by ImageJ. The 50% inhibitory concentrations (IC_50_) of PD-1 and CD45 on different targets were determined by fitting the dose response data in **B** using Graphpad Prism 5.0, or estimated based on the dose response plots if the inhibition was incomplete even at the highest PD-1 or CD45 concentration (summarized in **Table S2**). Error bars: S.D., N = 3.

To better understand the basis of the PD-1–Shp2 sensitivity to CD28, we deconstructed the reconstitution system into its individual modules (**fig. S11**). These experiments revealed that Shp2 alone dephosphorylates CD3ζ and CD28 with similar activities, but that Lck has a six-fold higher *k*_cat_ for CD3ζ over CD28 for phosphorylation (**fig. S11C-E**). These data suggest that the costimulatory receptor CD28 is a weak kinase substrate, which renders it more susceptible to PD-1–Shp2 inhibition in a kinase-phosphatase network. In summary, in reconstitution of components at physiological concentrations, CD28, and to a lesser extent Lck, are the major substrates for PD-1–Shp2 mediated dephosphorylation.

We next tested whether CD28 is the preferential target of PD-1 in intact T cells, as predicted from the biochemical reconstitution. For these studies, we used Jurkat T cells together with the Raji B cell line as an antigen presenting cell (APC), as this system has been widely used for studying TCR and CD28 signaling (*42*). Because these cells lack PD-1 and PD-L1, we lentivirally transduced PD-1 and PD-L1 into Jurkat and Raji respectively, obtaining PD-1+ Jurkat that express ∼40 molecules per µm^2^ PD-1 (**Table S1**) and Raji that express ∼86 molecules per µm^2^ PD-L1 (designated as PD-L1^High^, **Fig. 4A**). To titrate the strength of PD-L1–PD-1 signaling, the PD-1–expressing Jurkat T cells were incubated with different ratios of PD-L1^High^/PD-L1^Negative^ Raji B cells; since a T cell can interact with multiple APCs, this mixture of APCs might be expected to modulate the PD-1 response. Two minutes after APC and T cell contact, CD28 phosphorylation decreased as a function of the percentage of PD-L1^High^ cells (**Fig. 4B,C**, *t* = 2 min). In contrast, no or significantly less dephosphorylation was observed for ZAP70 and CD3ζ, respectively. Interestingly, PD-L1–PD-1 inhibitory effect was transient, with far less dephosphorylation detected at 10 min (**Fig. 4B,C**, *t* = 10 min), perhaps reflecting the feedback loop described *in vitro* (**Fig. 2H**) that enables recruited Shp2 to dephosphorylate PD-1 and thereby quenching the inhibitory signal. We next tested these results using a Raji B cell line that expresses lower levels of PD-L1 (∼16 molecules per µm^2^, designated PD-L1^Low^, **fig. S12A**), a density similar to that found in tumor infiltrating macrophages and tumor cells (**Table S3**). Using this lower-expressing APC line alone, we still detected a transient dephosphorylation of CD28 with little to no effect on TCR signaling components (**fig. S12B,C**, *t* = 2 min).

**Figure. 4.**
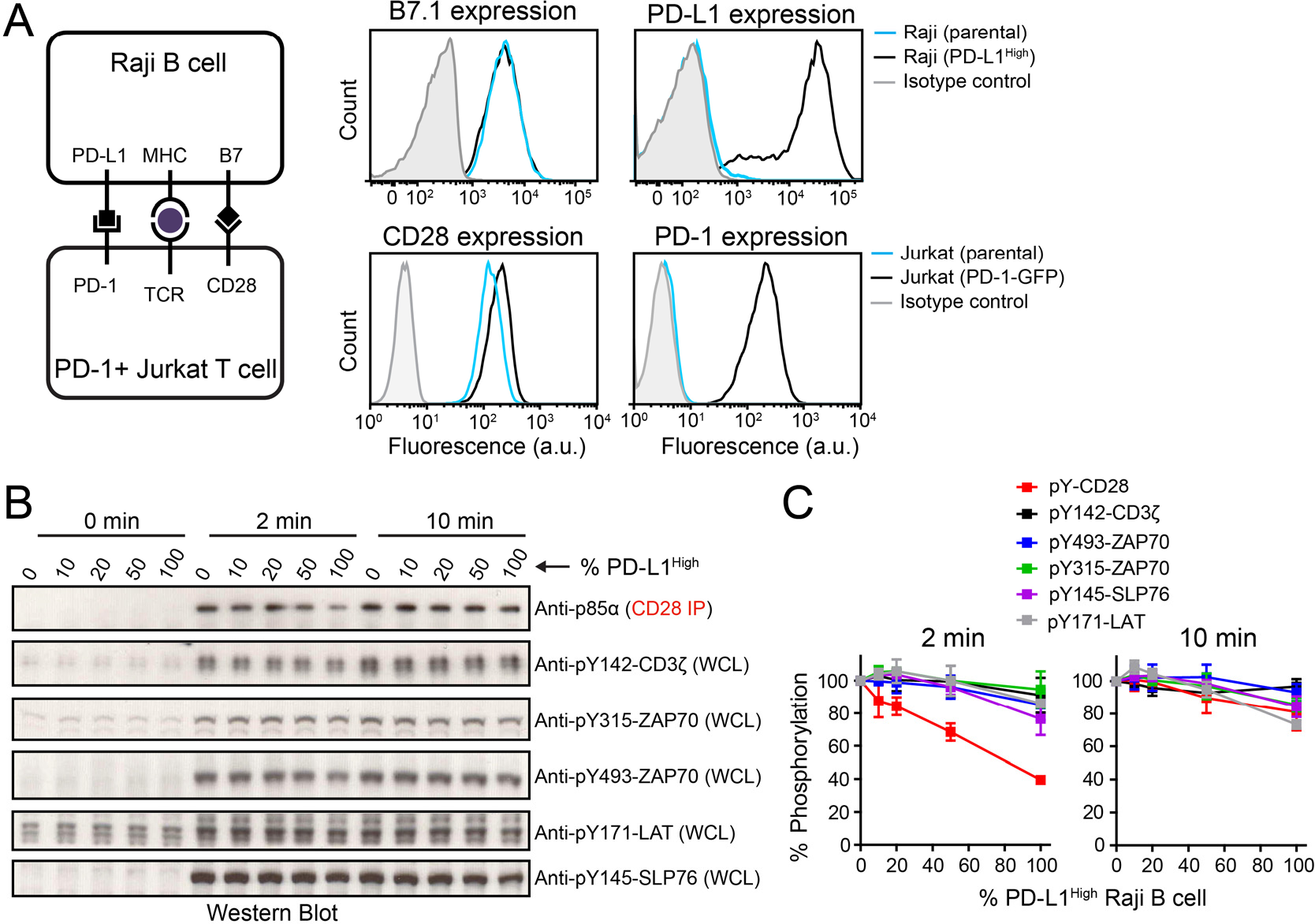
Intact cell assays confirmed CD28 as the preferential target of PD-1 mediated inhibition. (**A**) (*Left*) Cartoon (adapted from reference (*42*)) illustrating an intact cell assay in which CD28+, PD-1 transduced Jurkat T cells were stimulated with B7.1+, PD-L1 transduced (PD-L1^High^) Raji B cells preloaded with antigen. (*Right*) FACS histograms showing the expression of B7.1 and PD-L1 in parental or PD-L1^High^ Raji, and expression of CD28 and PD-1 in parental or PD-1 transduced Jurkat. **B**) A representative experiment of western blots showing the phosphorylation of CD28 and TCR signaling components in Jurkat T cells in response to PD-L1 titration on antigen-presenting Raji B cells; the time after the initial contact of the two cell populations is indicated (see **Methods**). Different ratios of PD-L1 ^High^ cells and PD-L1^Negative^ Raji B (both containing pMHC and B7.1) were used to vary the PD-L1 stimulation to the Jurkat cells. Each condition contained identical number of Raji B cells (Raji to Jurkat ratio = 0.75). The phosphorylation states of CD3ζ, ZAP70 and LAT were immunoblotted with phospho-specific antibodies. Due to lack of CD28 specific phosphotyrosine antibodies, CD28 was immunoprecipitated and co-immunoprecipitating p85α was measured by immunoblot, which is dependent upon the CD28 phosphorylation state (**C**) Quantification of phosphorylation data incorporating results from two independent experiments (mean and S.D., N = 3).

Taken together, results obtained from both membrane reconstitution and intact cell assays demonstrate that PD-1–Shp2 strongly favors dephosphorylation of the costimulatory receptor CD28 over TCR (**fig. S13**). Even at a relatively low levels of PD-L1 or PD-1, CD28 signaling remained selectively inhibited in both assays. At higher PD-L1 levels, we also observed some dephosphorylation of TCR components such as SLP76 and ZAP70. Although an effect on TCR signaling is in agreement with previous reports (*20-22*), the degree of TCR dephosphorylation was consistently much weaker than for CD28. To date, the relative propensities of these targets for regulation by PD-1 had not been directly or quantitatively compared.

The unexpected preference for inhibition of costimulatory receptor signaling may have interesting implications for cancer immunology and immunotherapy. Although costimulation via CD28 is most often associated with the priming of naïve T cells, there is increasing evidence that it may play a role at later stages of T cell immunity in cancer and in chronic virus infections. Indeed, recent studies have demonstrated that the ability of anti-PD-L1/PD-1 to rescue anti-viral (LCMV) T cell responses, as well as anti-tumor responses, depends on CD28 expression by T cells (Kamphorst et al., co-submitted). Blockade of B7.1/B.2 binding to CD28 also completely eliminated the ability of anti-PD-L1/PD-1 to rescue T cells from, or to prevent, exhaustion. These *in vivo* observations are entirely consistent with expectations from our results, namely that PD-1 exerts its primary effect by regulating CD28 signaling. In at least a subset of human cancer patients, inhibition of T cell immunity is associated with the upregulation of PD-L1 in the tumor bed in response to the release of IFNγ (*2, 6, 15, 16*). However, expression of PD-L1 by tumor infiltrating immune cells can be independently and possibly more predictive of clinical response than expression by the tumor cells themselves (*43*). The infiltrating cells including lymphocytes, monocytic cells, and dendritic cells, all express CD28 ligands while generally tumor cells do not. If the primary target of PD-1 signaling regulation is through CD28 or another costimulatory molecule, then the therapeutic effect is likely to reflect re-activation of costimulatory molecule signaling on T effector cells rather than (or at least in addition to) TCR signaling. Conceivably, costimulation is required to expand tumor antigen-specific early memory T cells, a process controlled intratumorally by B7.1+ APCs. Indeed, recent LCMV experiments have implicated an early memory population as being the targets for expansion by anti-PD-L1/PD-1 (*44, 45*). These findings strongly suggest that a role for costimulatory molecules, in addition to CD28 (which can be down-regulated in tumor infiltrating lymphocytes), in anti-tumor immunity must be strongly considered.

## ACKNOWLEDGMENTS

We thank Haopeng Wang (Shanghai-Tech University) and Fui-Boon Kai (UCSF) for help with retrovirus transduction of primary T cells, Nico Stuurman for training in TIRF microscopy, John James (University of Cambridge) for providing the lentiviral transfer plasmid pHR-PD-L1-mCherry, Arthur Weiss (UCSF) for providing the retrovirus vectors pMSCV and pCL-Eco, and Jon Ditlev (Michael Rosen’s lab, UT Southwestern Medical Center) for providing His_8_-LAT. We acknowledge members of the Vale lab for comments and discussions. R.D.V. is an investigator of the Howard Hughes Medical Institute.

